# Cis-element acquisition converts pervasive transcripts into translated mRNAs

**DOI:** 10.64898/2026.09.02.749023

**Authors:** Rongjia Zhang, Charles J. David

**Affiliations:** School of Medicine, Tsinghua University, Beijing, China; Peking University–Tsinghua Center for Life Sciences(CLS), Academy for Advanced Interdisciplinary Studies, Peking University, Beijing, China

**Keywords:** pervasive transcription, enhancer RNA, PROMPT, nuclear export, polyadenylation, de novo gene birth, mRNA fate

## Abstract

The mammalian genome is pervasively transcribed into thousands of unstable non-coding RNAs, including promoter-upstream transcripts (PROMPTs) and enhancer RNAs (eRNAs). These pervasive transcripts are degraded by the nuclear exosome within minutes of synthesis and are never translated. Coding and non-coding transcripts differ systematically in their content of splicing and polyadenylation signals, yet whether these cis-elements are sufficient to redirect a pervasive transcript toward an mRNA-like fate has not been tested at endogenous loci. Here, using CRISPR-Cas9 knock-in at three endogenous non-coding loci pervasively transcribed in murine cancer cells, we show that appending a splicing signal together with a polyadenylation signal is sufficient to render the resident transcript stable, polyadenylated, exported, and translated into full-length protein. Through quantitative analysis of isogenic single-cell clones, we find that the polyadenylation signal acts principally on transcript stability, extending half-life from ∼20 minutes to ∼12 hours, whereas splicing acts principally on nuclear export, with splicing efficiency quantitatively predicting both export and protein output. Remarkably, the untranslated regions of naturally intronless genes substitute for splicing, conferring coding capacity predominantly by enhancing export rather than stability. A genome-wide CRISPR screen on the intronless-gene reporter nominates nuclear export, 3′-end processing, and translation factors including NXF1, PABPN1, and EIF3G as required for conversion. Our results demonstrate that the acquisition of defined cis-elements is sufficient to convert pervasive transcripts into mRNAs, with nuclear export as the convergent rate-limiting step, providing an experimental reconstitution of a route by which non-coding loci may be recruited into the coding genome.

## Introduction

RNA polymerase II transcribes far more of the eukaryotic genome than is accounted for by protein-coding genes^1^. Much of this output is pervasive and non-coding: PROMPTs arise divergently from active promoters^2^, and eRNAs are transcribed bidirectionally from enhancers^3,4^. Unlike mRNAs, these transcripts are typically unspliced, non-polyadenylated, retained in the nucleus, and degraded within minutes of synthesis by the nuclear exosome and its associated machinery^5, 6^. This rapid turnover has made pervasive transcripts appear inert as protein-coding material, despite being produced abundantly across the genome.

A key molecular distinction separates the two transcript classes: protein-coding genes usually carry canonical 5′ splice, 3′ splice, and polyadenylation signals, whereas most pervasive-transcript loci lack them. The polyadenylation signal directs 3′ cleavage and polyadenylation and protects the transcript from exonucleolytic decay^7^, while splicing signals couple transcription to nuclear export through recruitment of the TREX complex and deposition of the exon-junction complex^8^. These correlations underlie a model of de novo gene birth, in which mutations that introduce splicing and polyadenylation signals into a non-coding locus allow its transcript to be exported and translated, exposing a previously cryptic peptide to selection^9^. Consistent with this model, comparative and population-genetic analyses identify nuclear export as the selectively constrained boundary that distinguishes mRNAs from lncRNAs during gene birth^9^. What has been missing is a direct test of sufficiency: whether installing these elements at an endogenous pervasive-transcript locus is enough to drive the transition.

Here we test this directly. By knocking defined cis-elements into three endogenous pervasive-transcript loci, including a PROMPT and two enhancers, and analyzing isogenic single-cell clones, we show that splicing plus polyadenylation signals are sufficient to convert a pervasive transcript into a stable, exported, translated mRNA. In the absence of splicing, we find that untranslated regions from intronless genes can also confer these properties. We identify enhanced nuclear export as the step through which these distinct sets of elemetns converge. Beyond the evolutionary question, the principle that a non-coding regulatory element can be rendered protein-coding carries practical utility: enhancers are numerous and exquisitely cell-type-specific, and recent work has begun to exploit appended processing signals to convert enhancers into compact, tissue-specific expression cassettes^10^. Defining which elements are sufficient, and the step at which they act, is therefore of value on both fronts.

## Results

### Splicing and polyadenylation signals render pervasive transcripts translated mRNAs

We studied three endogenous non-coding loci selected for their exosome-sensitive, non-coding transcription: E2, a PROMPT arising upstream of *Aqp5*; and R1 and J1, enhancer RNAs near *Tnfaip3* and *Bcl3*, respectively. All three were defined by transcripts that accumulated upon depletion of the exosome catalytic subunit DIS3 and of the PAF1 complex, confirming their identity as exosome-targeted pervasive transcripts (Figure 1A). R1 and J1 were marked by H3K4 monomethylation (H3K4me1) and H3K27 acetylation (H3K27ac), consistent with their status as enhancers. Using CRISPR-Cas9, we knocked a GFP reporter into each locus in one of three configurations: GFP alone; GFP followed by a polyadenylation signal (a 122-bp SV40 element bearing the conserved AATAAA hexamer); or GFP preceded by an exon–intron–exon (EIE) splicing cassette derived from internal, constitutively spliced exons from the *Arfgap2* gene with or without the polyadenylation signal (Figure 1B).

**Figure 1.**
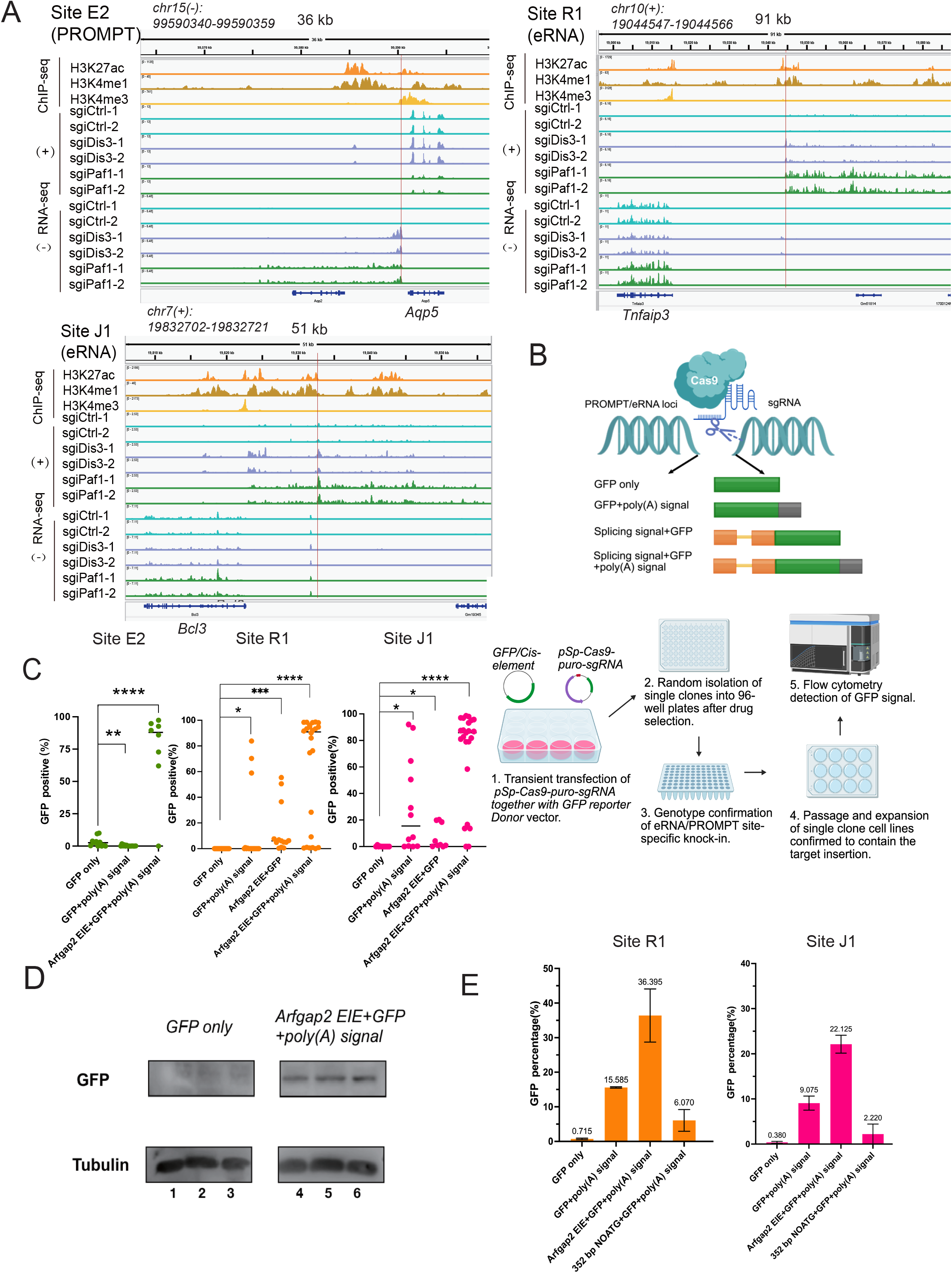
Splicing and polyadenylation signals render pervasive transcripts translated mRNAs. (A) Three non-coding loci—the E2 PROMPT (upstream of *Aqp5*) and the R1 (*Tnfaip3*) and J1 (*Bcl3*) enhancer RNAs—shown with ChIP-seq (H3K4me1, H3K4me3, H3K27ac) and strand-specific RNA-seq under control, *Dis3*, and *Paf1* knockout. Red lines indicate sgRNA target sites. (B) Schematic of the GFP reporter knock-in strategy and the splicing (EIE) and polyadenylation cis-elements. (C) Single-clone flow cytometry after knock-in of the indicated reporter constructs at the three loci. (D) Western blot for GFP in single clones: lanes 1–3, GFP-only (∼0% GFP); lanes 4–6, *Arfgap2* EIE+GFP+poly(A) (>90% GFP); α-tubulin, loading control. *\*P < 0.05, **P < 0.01, ***P < 0.001, ****P < 0.0001*. (E) Cell group flow cytometry analysis after transfection of indicated donor vectors at eRNA loci R1 and J1.

To avoid confounding effects of random integration and copy number, we isolated, genotyped, and propagated single-cell clones with verified on-target knock-in, and quantified GFP across multiple (10-15) independent clones per construct without pre-selection (Figure 1B). GFP alone yielded essentially no protein, and the polyadenylation signal alone produced only a modest increase. In contrast, combining the splicing cassette with the polyadenylation signal drove robust GFP expression at all three loci (median ∼90% GFP-positive at R1 and ∼85% at J1, with comparable conversion at the E2 PROMPT)(Figure 1C). Western blotting confirmed accumulation of full-length GFP protein in converted clones but not in GFP-only clones (Figure 1D). To rule out changes in the length of the 5’ UTR of the GFP coding sequence accounting for the EIE-induced increase in protein production, we made constructs containing ATG-free bacterial-derived sequences equivalent in length to the EIE cassette (352 bp). This resulted in no increase in GFP production (Figure 1E), suggesting that splicing signals, and not lengthening accounted for the increased protein production. Thus, the addition of splicing and polyadenylation signals is sufficient to convert an endogenous pervasive transcript, whether a PROMPT or an eRNA, into a translated mRNA.

### Polyadenylation confers stability, splicing confers export, and splicing efficiency sets output

To dissect the contribution of each element, focusing on the eRNA sites R1 and J1, we analyzed the converted single-cell clones across four readouts. First, measuring polyadenylation status as the ratio of oligo(dT)- to random-hexamer-primed reverse transcription, we found that, as expected, polyadenylation-signal–containing constructs were markedly more polyadenylated than GFP alone across the eRNA loci, whereas the EIE cassette without a polyadenylation signal was not (Figure 2A). Second, the polyadenylation signal extended transcript half-life from ∼20 minutes for GFP alone to ∼12 hours (Figure 2B). Third, in line with the stabilization, steady-state transcript abundance rose with the polyadenylation signal and further with the combined EIE-plus-polyadenylation construct at the eRNA loci (Figure 2C). Having established stabilization, we examined the export of transcripts from the eRNA loci. Only the full splicing-plus-polyadenylation configuration substantially increased the cytoplasmic export ratio; neither the polyadenylation signal alone nor the EIE cassette lacking a polyadenylation signal was sufficient to drive export (Figure 2D). To establish causality of the splicing signals, we introduced targeted mutations: disrupting the 3′ splice site, or deleting the polyadenylation signal, strongly reduced GFP, while 5′ splice-site mutation and intron or exon manipulations produced graded effects (Figure 2E). We were surprised that some mutations expected to abolish splicing still permitted GFP production in some clones. We therefore examined whether the mutations indeed abolished splicing using RT-PCR. Surprisingly, some clones continued to produce considerable amounts of spliced mRNA, despite the 3’ or 5’ SS mutation, suggesting the use of cryptic splice sites (Figure 2F). Strikingly, across mutant single clones, splicing efficiency quantitatively predicted both GFP output (Pearson r = 0.76 for 5′ splice-site and 0.84 for 3′ splice-site mutant series) and export efficiency (r = 0.97 and 0.85, respectively) (Figure 2G, 2H). Together these data indicate that the polyadenylation signal acts chiefly on stability, the splicing signal acts chiefly on export, and the efficiency of splicing-coupled export sets the level of protein output.

**Figure 2.**
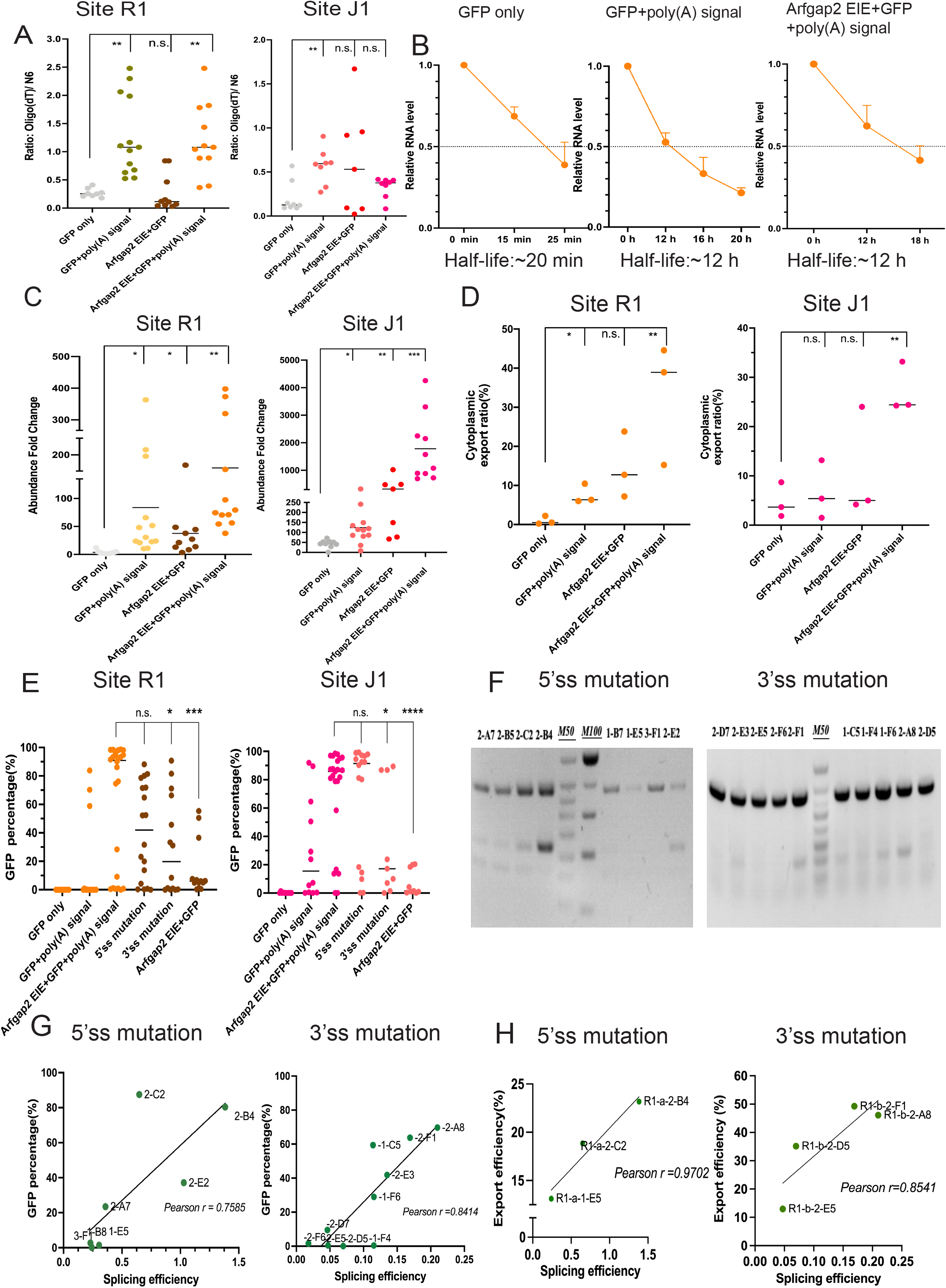
Polyadenylation confers stability, splicing confers export, and splicing efficiency sets output. (A) Polyadenylation status (oligo(dT)/N6 ratio) of single clones at the three loci. (B) RNA half-life of three representative clones at R1. (C) Steady-state reporter abundance at the three loci. (D) Cytoplasmic export ratio of single clones at the two eRNA loci (R1 and J1). (E) Single-clone GFP for splice-site mutants, element deletions, and polyadenylation-signal deletion at R1 and J1.(F) RT-PCR analysis of 5′ and 3′ splice-site mutant single clones. (G) Correlation of splicing efficiency with GFP for 5′ and 3′ splice-site mutant clones. (H) Correlation of export efficiency with splicing efficiency for the same clones. *\*P < 0.05, **P < 0.01, ***P < 0.001, ****P < 0.0001*; n.s., not significant.

### Intronless-gene UTRs convert enhancer transcripts via export, independently of splicing

Reasoning that splicing might act largely by licensing nuclear export, we asked whether export could instead be conferred without any splicing events. Naturally intronless protein-coding genes export their mRNAs despite never being spliced, through cis-acting elements within their transcribed sequences. We therefore replaced the splicing cassette with the 5′ and 3′ untranslated regions (UTRs) of three intronless genes—*Sephs2, H1f0*, and *Ddx28*—flanking GFP, retaining the polyadenylation signal (Figure 3A). Each UTR pair conferred GFP expression at both R1 and J1, comparable to the splicing-based cassette (Figure 3B). In single-cell clones, the *Sephs2*-UTR reporter did not increase transcript abundance relative to GFP-plus-polyadenylation (not significant) but significantly increased the cytoplasmic export ratio (Figure 3C, 3D). Thus, intronless-gene UTRs drive conversion predominantly by enhancing nuclear export rather than transcript stability—mechanistically distinct from, but functionally convergent with, the splicing route.

**Figure 3.**
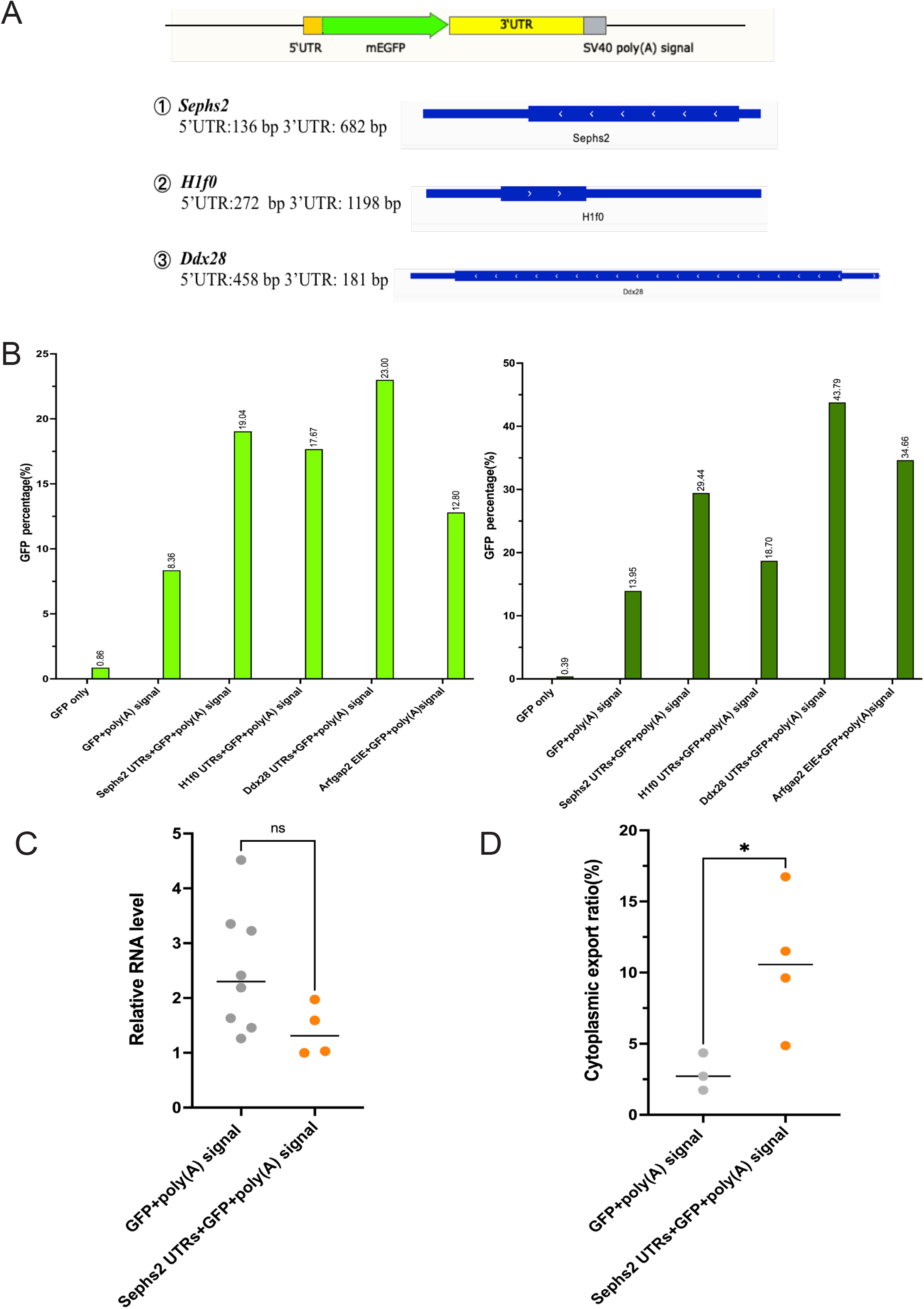
Intronless-gene UTRs convert enhancer transcripts via export, independently of splicing. (A) Schematic of the intronless-gene 5′UTR–mEGFP–3′UTR–poly(A) reporter; UTRs from *Sephs2, H1f0*, and *Ddx28*. (B) Flow cytometry of the UTR reporters at R1 and J1 (negative control, GFP only; positive control, *Arfgap2* EIE+GFP+poly(A)). (C, D) Comparative RNA abundance (C) and cytoplasmic export ratio (D) of *Sephs2*-UTR versus GFP+poly(A) single clones at R1. *P < 0.05; n.s., not significant.

### A genome-wide CRISPR screen identifies conversion factors

To identify trans-acting factors required for UTR-mediated conversion, we performed a genome-wide CRISPR knockout screen in a clonal line carrying the *Sephs2*-UTR–GFP–polyadenylation reporter at R1 together with doxycycline-inducible Cas9 (R1-*Sephs2*-Cas9-2). Following transduction with the mouse Brie knockout library, selection, and Cas9 induction, we sorted GFP-low (P4, ∼7%) and GFP-high (P5, ∼80%) populations and identified sgRNAs enriched in the GFP-low fraction by MAGeCK^11^ (Figure 4A, 4B). Top hits included the principal mRNA export receptor NXF1, the nuclear poly(A)-binding protein PABPN1, and the translation-initiation factor EIF3G, alongside numerous ribosomal proteins and several canonical histone genes (Figure 4C). Surprisingly, the splicing factor SF3B5 was also a strong hit, suggesting a splicing-independent role for this factor in the export of intronless genes. Individual sgRNA validation in the reporter line confirmed that knockout of NXF1, PABPN1, SF3B5, EIF3G, and ribosomal proteins reduced GFP relative to non-targeting controls (Figure 4D, 4E). We interpret NXF1, PABPN1, and EIF3G as on-pathway factors consistent with an export–processing–translation route. The enrichment of ribosomal proteins and canonical histones, by contrast, most likely reflects general essentiality in a viability- and expression-coupled sort rather than a dedicated role in fate conversion; distinguishing specific from general requirements will need a parallel screen on a constitutively expressed reporter.

**Figure 4.**
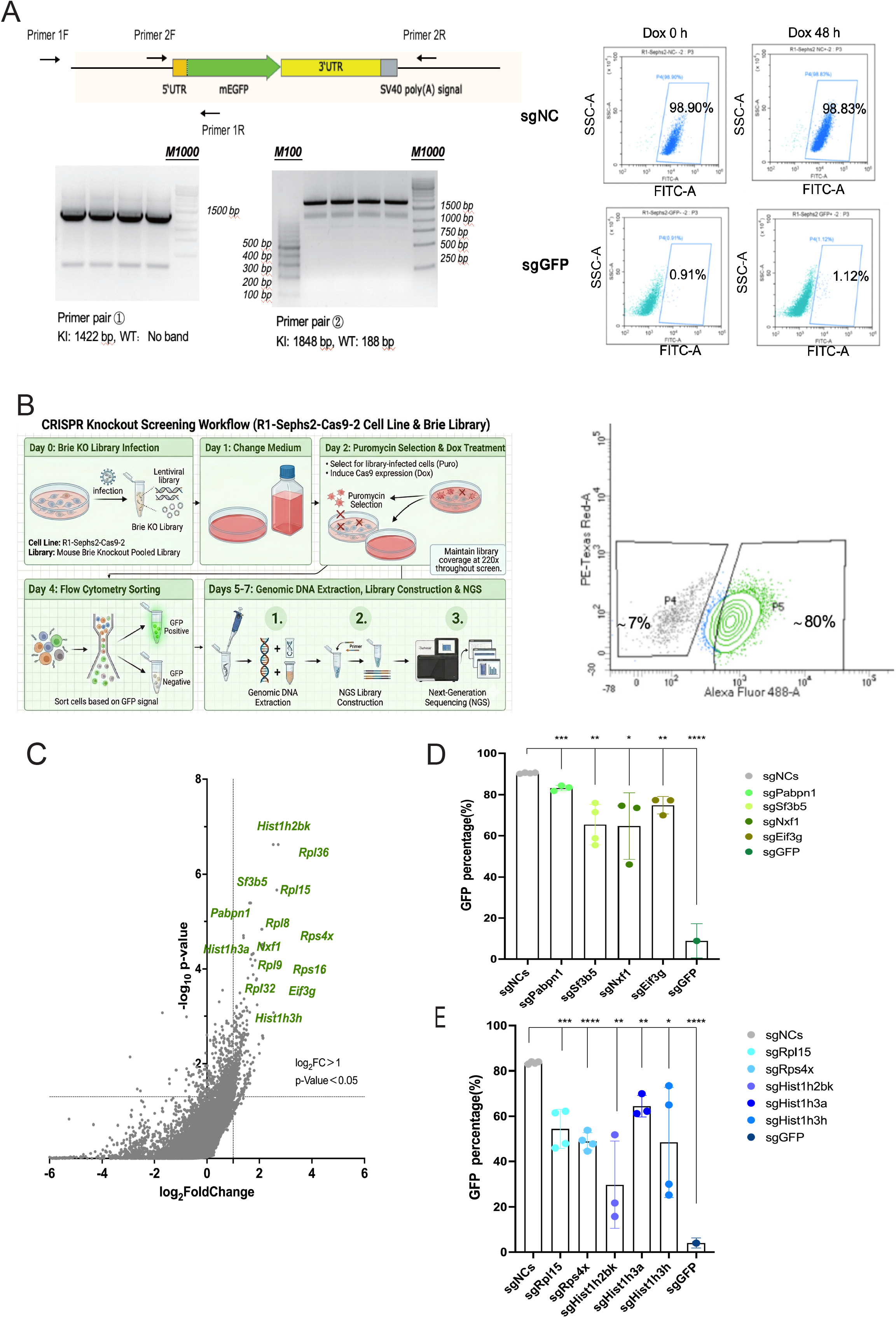
A genome-wide CRISPR screen nominates conversion factors. (A) Genotyping and flow cytometry test of screening cell line R1-*Sephs2*-Cas9-2. (B) Screen workflow and GFP-low/GFP-high sorting gates (P4, ∼7%; P5, ∼80%) in the R1-*Sephs2*-Cas9-2 line. (C) MAGeCK analysis of sgRNAs enriched in the GFP-low fraction (log2FC > 1, P < 0.05). (D, E) Individual-sgRNA validation in the reporter line (sgNC, non-targeting; sg*GFP*, positive control). n = 3 independent experiments. *\*P < 0.05, **P < 0.01, ***P < 0.001, ****P < 0.0001*.

## Discussion

These experiments provide a direct test of cis-element sufficiency for RNA fate. At endogenous PROMPT and enhancer loci whose transcripts are normally degraded within minutes, the addition of a splicing signal and a polyadenylation signal is sufficient to produce a stable, exported, and translated mRNA. Quantitative dissection separates the contributions of each: polyadenylation stabilizes the transcript, splicing licenses its export, and the efficiency of splicing-coupled export rather than stability or abundance sets the amount of protein made. The convergence of a second, splicing-independent route built from intronless-gene UTRs on the same export step identifies nuclear export as the rate-limiting node in the conversion of a pervasive transcript into an mRNA.

This result offers an experimental reconstitution of the central step in the “stowaway” model of de novo gene birth. Where comparative and population-genetic analyses have inferred that the acquisition of splicing- and export-promoting elements marks the boundary a nascent transcript must cross to become mRNA-like, our knock-ins show directly that installing such elements at a non-coding locus is sufficient to drive that transition. That two mechanistically distinct element sets, spliceosomal and intronless-gene UTRs, both act through export reinforces the view that escaping nuclear retention and exosomal decay, rather than acquiring a stable poly(A) tail per se, is the decisive event in recruiting a pervasive transcript into the translated genome.

The same principle has begun to be exploited for synthetic biology, where appending processing signals can turn enhancers into compact, cell-type-specific expression cassettes for reporter and therapeutic genes. Our study is complementary and distinct: rather than optimizing expression from a neutral landing site, we test conversion at the endogenous non-coding loci themselves, dissect the quantitative contribution of each element, and use an unbiased genetic screen to nominate the trans-acting machinery. NXF1, PABPN1, and EIF3G, the factors recovered from the screen, map onto the export, 3′-end-processing, and translation steps that the cis-element analysis predicts.

Our conclusions are constrained by several limitations. The experiments were performed in a single cell line, and confirming generality will require additional cell types and, ideally, an in vivo context. Our reporter supplies its own open reading frame, so we demonstrate that the cis-elements are sufficient to confer an mRNA-like fate on a transcript arising from a non-coding locus, rather than translation of the native enhancer sequence. The conversion itself was demonstrated at three loci spanning both the PROMPT and enhancer classes, supporting the generality of the mechanism across pervasive transcripts; the detailed export and splicing-mutation dissection, however, was performed at the two eRNA loci, and confirming that the same export-centric logic operates at PROMPT loci will benefit from the corresponding mechanistic measurements. Notwithstanding these caveats, the demonstration that defined cis-elements are sufficient to convert an endogenous pervasive transcript into a translated mRNA, and that nuclear export is the convergent step, clarifies both how the coding genome may be expanded over evolution and how non-coding transcription might be redirected by design.

## Materials and Methods

### Cell culture

All experiments were performed in mouse pancreatic cancer cell line 806 (*Pdx1-cre; LSL-Kras*^*G12D*^; *Ink4a*^*fl/fl*^; *Smad4*^*fl/f*l^). Cells were cultured in Dulbecco’s Modified Eagle Medium (DMEM) supplemented with 10% fetal bovine serum and 1% penicillin/streptomycin at 37°C and 5% CO2.

### CRISPR-Cas9 knock-in and single-clone isolation

sgRNAs targeting the R1 and J1 loci were cloned into pSpCas9(BB)-2A-Puro (PX459) and co-transfected with the corresponding GFP-reporter donor plasmid. Following transfection, cells were seeded at limiting dilution; single-cell clones were expanded and genotyped by PCR across both homology arms to confirm on-target integration. For each construct and KI location, 10-15 verified clones were analyzed without GFP pre-selection.

### Reporter constructs

The polyadenylation signal was a 122-bp SV40 late element containing the AATAAA hexamer. Splicing cassettes comprised an internal exon–intron–exon (EIE) unit from *Arfgap2*. Intronless-gene reporters carried the 5′ and 3′ UTRs of *Sephs2* (136/682 bp), *H1f0* (272/1198 bp), or *Ddx28* (458/181 bp) flanking mEGFP. Splice-site mutants and element deletions were generated by homology-directed recombination.

### Flow cytometry

GFP fluorescence was quantified on Cytoflex S (BECKMAN COULTER); Gating was set on non-transfected/parental controls. Data were analyzed in FlowJo (version 3.05478). For single-clone experiments, each point represents an independent clone.

### RNA half-life

Cells were seeded in 6-well plates 24 hrs before transcription inhibition. Transcription was blocked with (10 μg/mL); RNA was harvested at the indicated times and reporter transcript levels quantified by RT-qPCR relative to t = 0. Half-life was estimated by the time point at which relative RNA level decreased to 50%.

### Subcellular fractionation and export ratio

Cytoplasmic, nucleoplasmic, and chromatin fractions were separated as follow:Cells were first trypsinized and washed with ice cold PBS. Cell pellets were then lysed in an ice cold NP-40 homogenization buffer (10 mM Hepes-KOH [pH 7.6], 15 mM KCl, 2 mM EDTA, 0.15 mM spermine, 0.5 mM spermidine, 10% glycerol, 0.3M sucrose, 0.5% NP-40) and incubated 10 min on ice. A fraction of the lysate was used to prepare total RNA. NP-40 lysates were then layered on 30% cushion buffers (10 mM Hepes-KOH [pH 7.6], 15 mM KCl, 2 mM EDTA, 0.15 mM spermine, 0.5 mM spermidine, 10% glycerol, 0.87 M sucrose) in centrifugation tubes and spun 15 min at 3,500xg at 4°C. Isolated nuclei were resuspended in nuclear storage buffer (10 mM Hepes-KOH [pH 7.6], 100 mM KCl, 0.1 mM EDTA, 0.15 mM spermine, 0.5 mM spermidine, 10% glycerol). One volume of nuclei in nuclear storage buffer was then extracted with 1 volume of 2X NUN buffer (50 mM Hepes-KOH [pH 7.6] 0.6 M NaCl, 2% NP-40, 2M Urea) for 30 min on ice.

After centrifugation for 30 min at 21,000xg at 4°C, chromatin pellets were washed twice with 1X NUN buffer. The pellets were then thoroughly resuspended in TRizol until complete decompaction and then incubated 10 min at room temperature. Fractionation quality was monitored using the unspliced *Gapdh* intron as a nuclear marker. Cytoplasmic export ratio was calculated as cytoplasmic over total reporter transcript.

### Polyadenylation status

Reverse transcription was primed in parallel with oligo(dT) and random hexamers (N6); the oligo(dT)/N6 ratio of reporter transcript, measured by qPCR, served as an index of polyadenylation.

### Splicing efficiency

Spliced and unspliced reporter species were quantified by RT-PCR; splicing efficiency was defined as spliced over total. Correlations with GFP and export were assessed by Pearson correlation.

### Genome-wide CRISPR screen

The R1-*Sephs2*-Cas9-2 line was transduced with the mouse Brie knockout library at low MOI(∼0.20-0.25), maintaining ∼220× coverage. After puromycin selection and doxycycline-induced Cas9 expression, GFP-low and GFP-high populations were sorted; genomic DNA was extracted, sgRNA cassettes amplified, and libraries sequenced. Enrichment was analyzed with MAGeCK (version 0.5.9.2). Candidate sgRNAs were validated individually in the reporter line (n = 3 independent experiments).

### Western blot

Lysates were resolved by SDS-PAGE, transferred, and probed for GFP and α-tubulin (loading control).

#### Extended version

Cells used for western blot were lysed in 1xloading buffer (200mM Tris-HCl pH 6.8, 10% glycerol, 2% SDS, 0.1% bromophenol blue and 1% b-mercaptoethanol). Samples containing 1x10^6^ cells in each 100 μL loading buffer were boiled at 95^°^C for 5 min. Proteins were transferred to nitrocellulose filter membrane, blocked with 5% BSA, and incubated overnight with anti-GFP or anti-tubulin(loading control). HRP-conjugated secondary antibody was used as secondary antibody.

### Statistics

GraphPad Prism 9.0 and Excel software were used for statistical analysis. Statistical analysis between group comparisons was performed by unpaired Student’s t test. Significance levels are indicated as ∗, p value <0.05, ∗∗, p value <0.01, ∗∗∗, p value <0.001 and ∗∗∗∗, p value<0.0001. Please see figure legends for detailed information.

